# In situ cryo-electron tomography reveals filamentous actin within the microtubule lumen

**DOI:** 10.1101/844043

**Authors:** Danielle M Paul, Judith Mantell, Ufuk Borucu, Jennifer Coombs, Katherine J Surridge, John Squire, Paul Verkade, Mark P Dodding

## Abstract

Microtubules and filamentous (F-) actin engage in complex interactions to drive many cellular processes from subcellular organisation to cell division and migration. This is thought to be largely controlled by proteins that interface between the two structurally distinct cytoskeletal components. Here, we use cryo-electron tomography to demonstrate that the microtubule lumen can be occupied by extended segments of F-actin in small-molecule induced, microtubule-based cellular projections. We uncover an unexpected versatility in cytoskeletal form that may prompt a significant development of our current models of cellular architecture and offer a new experimental approach for the *in-situ* study of microtubule structure and contents.

## Introduction

Understanding of the lumenal contents of cytoplasmic microtubules has been classically driven by electron microscopy based analysis in cells and tissue ^1-3^, and most recently by high resolution cryo-electron microscopy (cryo-EM) ^4-8^. Contents of cytoplasmic microtubules are thought to be restricted to globular proteins such as tubulin modifying enzymes ^9^, although higher order structures have been observed in sperm flagella microtubules and cilia ^10,11^. Unequivocal identification of lumenal proteins in a cellular context remains a challenge as the dimensions of the microtubule may confound super-resolution fluorescence imaging techniques and there are potential issues with antibody epitope accessibility in this confined environment. Cryo-EM solves the challenge of spatial resolution but can be limited by the technical requirement for very thin samples ^12^ and so studies have mainly focused on microtubules at the cell periphery or within neuronal processes ^4-8^. We recently identified a compound, 3,5-dibromo-N′-[2,5-dimethyl-1-(3-nitrophenyl)-1H-pyrrol-3-yl]methylene}-4-hydroxybenzohydrazide (named ‘kinesore’), that targets the motor protein kinesin-1, promoting extensive remodelling of the microtubule network ^13^. Kinesore treatment results in dynamic looping and bundling of microtubules within the cytoplasm and their extrusion from the cell body as membrane bound projections. This renders microtubules from within the cell accessible for cryo-EM studies ^13,14^. Here we describe the discovery of filamentous actin (F-actin) inside the microtubule lumen through *in situ* cryo-electron tomography (Cryo-ET) analysis of these small molecule induced projections.

## Results

### Formation and ultrastructure of small-molecule induced microtubule-based projections

To begin ultrastuctural analysis of projections in their native state, HAP1 cells, incubated with a fluorescent membrane stain prior to kinesore treatment, were prepared for cyro Correlative Light Electron Microscopy (Cryo-CLEM). This allowed the unambiguous identification of projections using fluorescence microscopy that could be correlated with images from the electron microscope (Figure 1A). Consistent with our earlier immunofluorescence imaging study ^13^, abundant projections were composed of closely aligned microtubules and we also occasionally observed vesicular structures within swollen regions. Our previous live-imaging of GFP-tubulin expressing HeLa cells suggested that the projections were initially formed though the extrusion of microtubule loops ^13^. Live-imaging of SiR-tubulin labelled microtubules in the HAP1 cells used here confirmed this and showed that microtubules are progressively added through additional loop extrusion events (Extended Data Fig. 1 and Movie 1), providing a rationale for the formation of extended microtubule bundles. Consistent with this live-imaging, EM analysis of regions proximal to the cell body revealed a greater number of microtubules and distinct looped bundles (Extended Data Fig. 2). Within these structures, microtubules typically maintained a consistent spacing of between 10-25 nm (blue shading) although some were also observed to traverse bundles (yellow shading).

**Figure 1.**
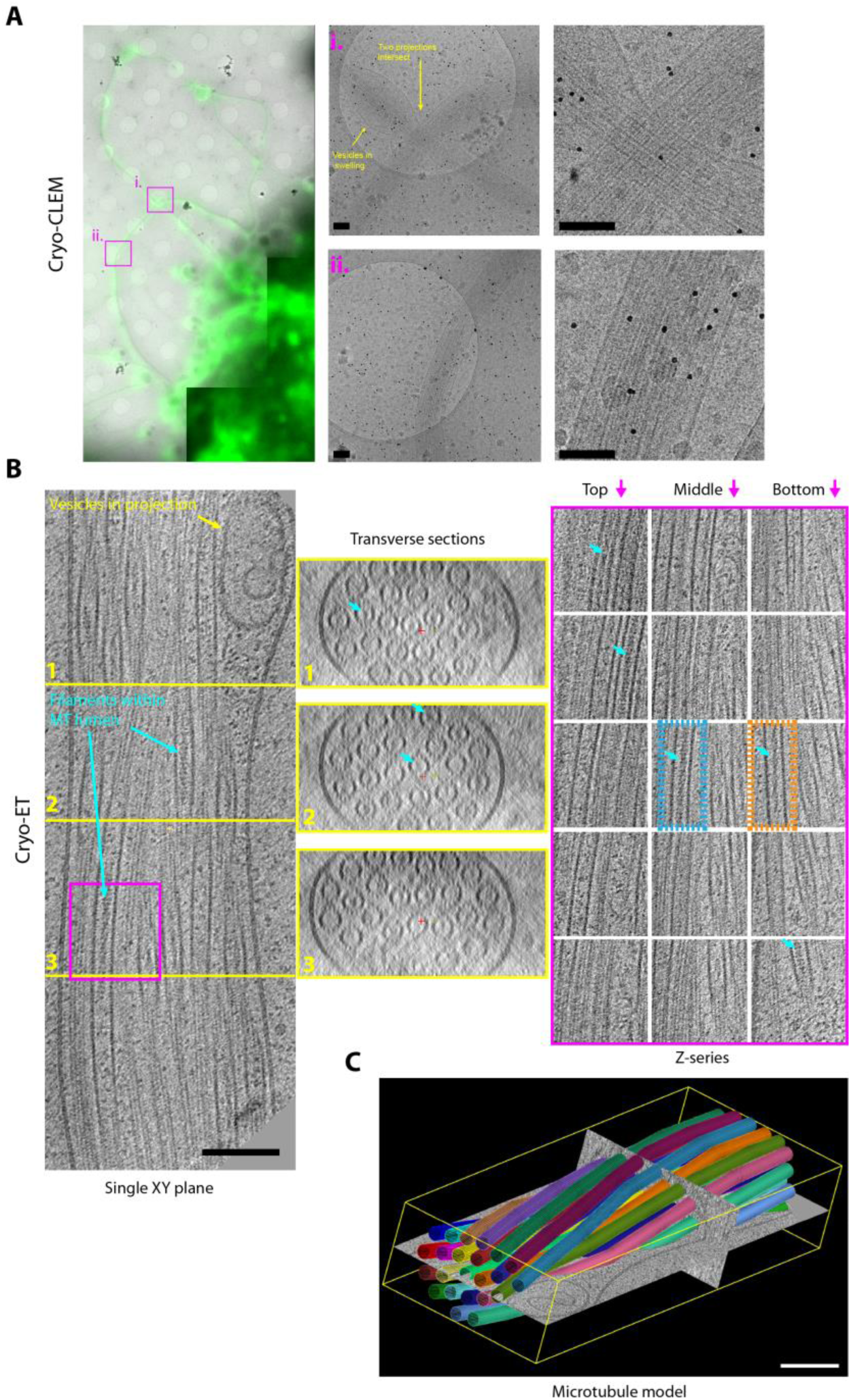
Cryo-EM reveals actin-like filaments within the microtubule lumen. HAP1 cells grown on Quantifoil gold EM grids were stained with Cell Mask green to identify plasma membrane and treated with kinesore (100µM) prior to plunge freezing and imaging using fluorescence and electron microscopy (A) Low magnification images shows fluorescence (green) and EM image (greys) overlayed of cells with projections emerging. Regions boxed i. and ii. are shown at higher magnification on the right. Black dots are 10nm gold fiducials. (B) An X-Y plane (left), a series of transverse sections (middle), and a Z-series (right) though a 3D reconstruction of 23-microtubule projection. Sections which bisect the microtubule lumen reveal a mixture of particles and filamentous material with apparent helical properties. Blue arrows highlight several lumenal filaments. Orange box regions highlight filaments of the Class I variety and blue box Class II as described in more detail in figure 2. (C) Shows a 3D model where microtubules are represented as coloured 24nm diameter cylinders segments (corresponds to Supplementary Movie 2). Microtubules are organised in a closely packed twisted membrane bound bundle with some vesicular structures in swellings at the periphery.

**Figure 2.**
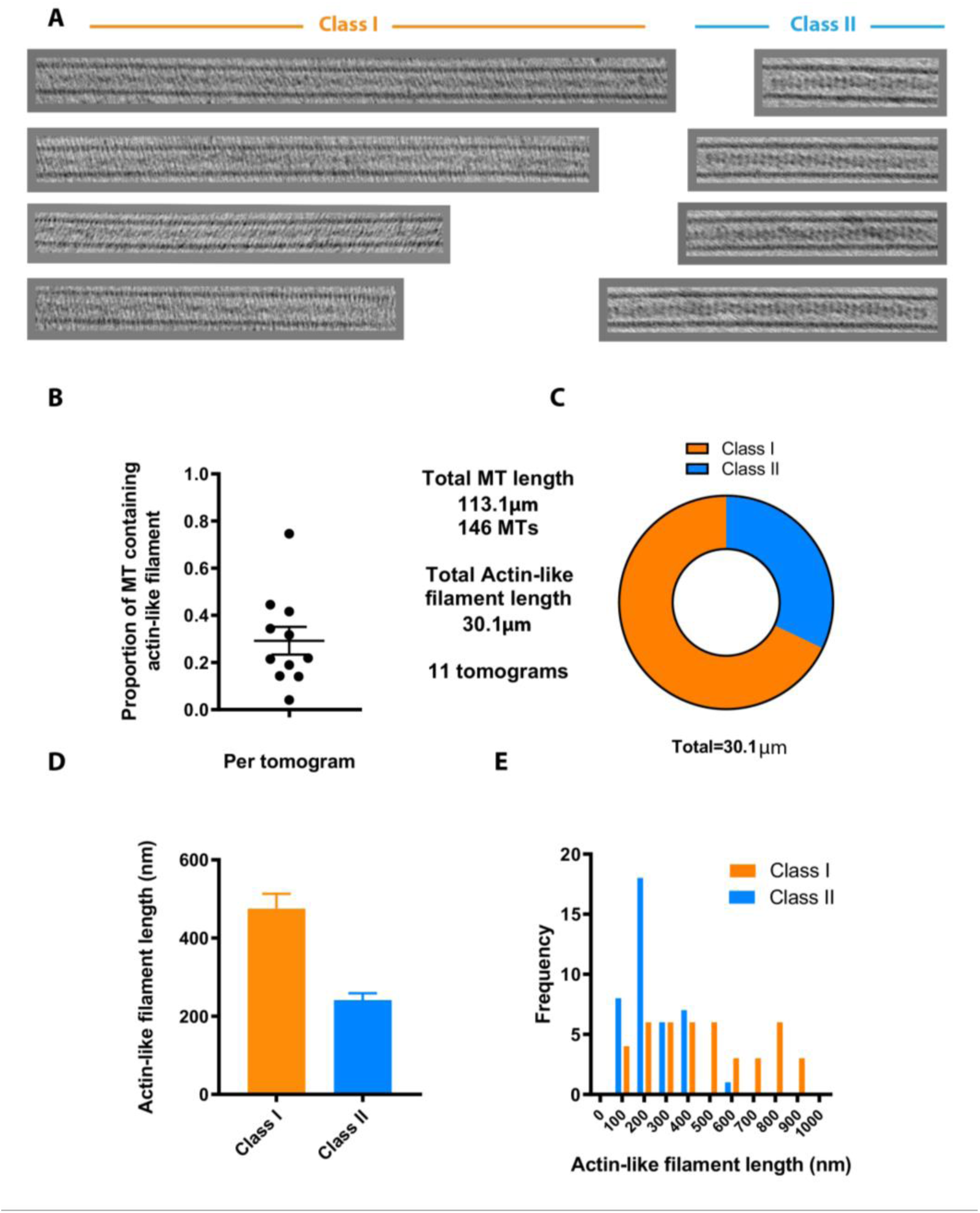
Abundance and classification of two types of actin-like filament. (A) Examples of lumenal filaments extracted from tomogram subvolumes, chosen from straight sections of microtubule so that lumenal filaments can be shown over an extended length without being obscured by surrounding microtubule lattice. Images show projected sections of central regions of microtubules containing actin-like filaments and are separated into two groups (Class I and Class II) based on their gross morphology. Class II filament were thicker and typically better defined (B) Graph shows the proportion of MT lumen occupied by actin-like filaments (of either class) in 11 tomograms. (C) Proportion of actin-like filament length classified as Class I or Class II from the dataset. (D) Graph showing average length of actin-like filaments (error bar s.e.m.) of either class and their frequency distribution (E).

### Cryo-ET reveals actin-like filaments with the lumens of extruded microtubules

Satisfied that we could confidently identify projections in the electron microscope, additional samples were analysed without the fluorescence imaging step (avoiding ice contamination) by 3D reconstruction of tomography tilt series (Figure 1 B,C). Transverse sections through projections revealed that they could contain variable numbers of microtubules (ranging from 4 to >30) (Figure 1B, Extended Data Fig. 3). A section of a 23 microtubule projection is shown in detail. Microtubules are organised in a twisted bundle and vesicles are excluded in swellings proximal to the limiting membrane. Both transverse sections and serial images of Z-sections show that the microtubules are intact (Figure 1C, Movie 2). Notably, these microtubule structures closely resemble those observed in *in vitro* EM studies of kinesin-driven microtubule-microtubule cross-linking ^15^. Surprisingly, within the microtubule lumens, as well as globular structures we could clearly observe extended filamentous density with helical character (blue arrows). Two examples of such filaments are boxed in orange and blue on the Z-section panels in Figure 1B and are also visible as density within the microtubule lumen of transverse sections (see also, Extended Data Fig. 3). The diameter (5-9 nm) and helical appearance of these lumenal filaments is consistent with that of F-actin ^16,17^, that at least in principle, could reside within the ≈ 15-16 nm diameter lumenal space ^18^.

To assess whether kinesore-induced projections do indeed contain actin, methanol fixed cells (to optimally preserve microtubules) were stained with antibodies against actin and tubulin. This revealed patches and puncta of actin along microtubules within the projections (Extended Data Fig. 4A). Actin antibodies may not discriminate between G and F-actin and so equivalent samples were prepared using paraformaldehyde fixation and phalloidin staining for F-actin. Under these fixation conditions, projections were less well preserved, with fragmentation in β-tubulin staining but actin patches were more prominent and appeared to bridge gaps in tubulin staining (Extended Data Fig. 4B). This raises the intriguing possibility that a population of F-actin may reside within the microtubule lumen that is refractory to traditional means of detection, possibly due to limited antibody epitope or small-molecule binding site accessibility.

**Figure 3.**
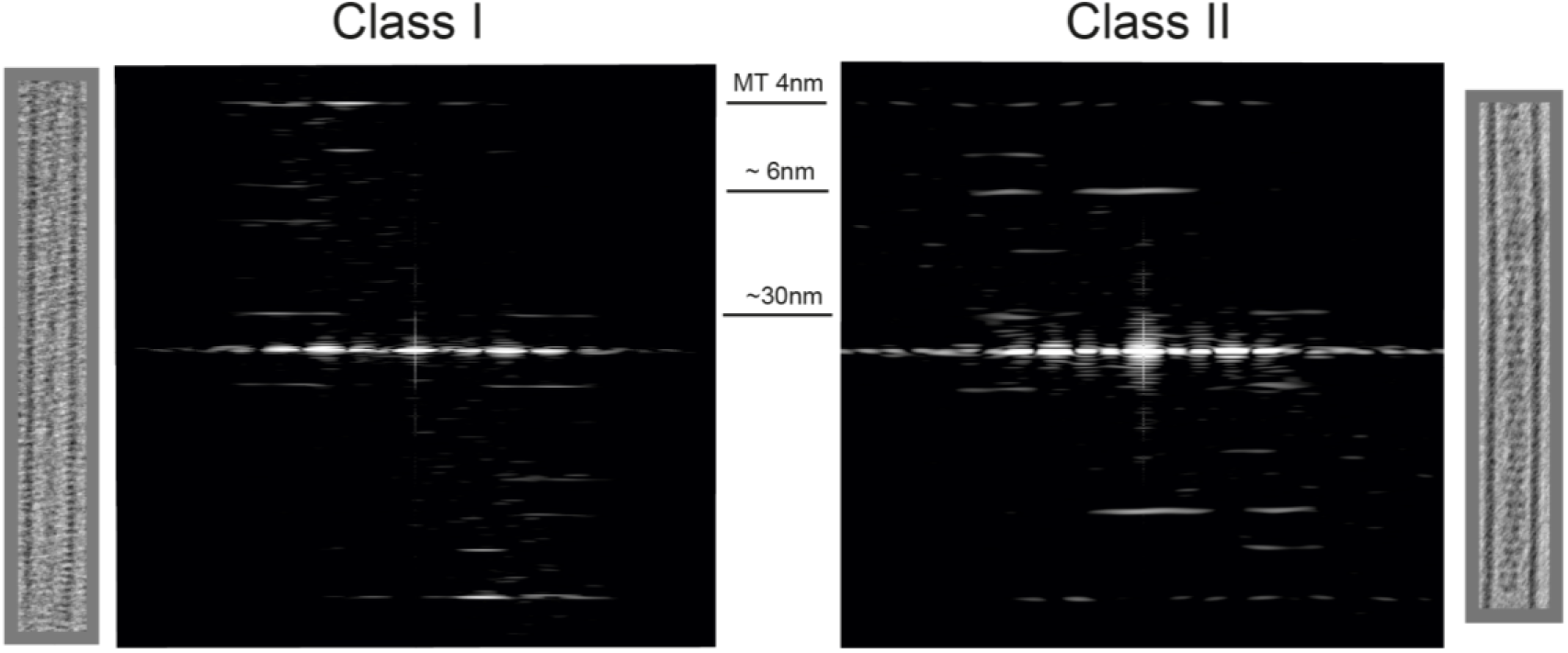
Analysis of layer line patterns of Class I and Class II filaments reveals typical actin repeating structures. Typical Class I (left) and Class II (right) luminal filaments are extracted as 3D sub-volumes from the tomographic reconstruction and shown as projected in Z. The corresponding layer line patterns are shown beside the projection. The characteristic 4nm tubulin layer line was visible and in all patterns. Parameters measured for more filaments are provided in Table 1. Class I filaments exhibited a clear reflection at 5.94 ±0.03nm the pitch of the genetic helix of actin and a reflection at 29.50± 0.80nm corresponding to the crossover spacing as measured in the individual images. Reflections were observed in the Class II filaments at similar positions to Class I 6.11 ±0.09 and 27.44 ±2.29nm, with an additional strong meridional reflection at 6.18±0.06nm (errors are S.D.). S.E.M. is provided in Table 1.

### Lumenal actin-like filaments have two distinct morphologies

A gross survey of the morphology of lumenal filaments resulted in their classification into two pools; Class I and Class II. Class I filaments are exemplified in the right orange box in Figure 1B Z-sections and class II by the left blue box. Further examples are shown in Figure 2A and Movie 3 shows a Z-series through an extended Class I filament containing microtubule. Class II filaments appeared slightly thicker and were typically better defined than Class I. Analysis of the tomograms in our dataset that provided clearest definition of lumenal contents (n=11), containing 146 microtubules with a total length of 113 µm revealed that 27% of the total lumenal length of microtubule was occupied by actin-like filaments of either class (Figure 2B). There was considerable variation between tomograms with 76% to 4% of the lumenal length occupied, with a mean of 29%. Of the total actin-like filament length, 68% was comprised of Class I and 32% by Class II filaments (Figure 2C). Although the frequency of Class I and Class II filaments were similar, Class I filaments were longer (average 475nm) than Class II (average 274nm) (Figure 2 D,E). Class I filament containing microtubules had average outer and lumenal diameters of 25.60 ± 0.59 nm and 16.80 ± 0.76 nm respectively. Class II filaments containing microtubules were typically slightly wider at 27.29 ± 0.58 nm (outer diameter) and 17.80 ± 0.85 nm (lumenal diameter) indicating that the presence of lumenal filaments either correlates with or modifies microtubule properties (Table 1).

**Table 1.** Measured parameters for individual Class I and Class II filaments. MT dimensions and layer line positions are listed for filaments of both classes. All data are presented in angstroms.

| **Class I** | **Crossover spacing** | **Turn layer-line** | **MT outer diameter** | **MT lumen diameter** |
| --- | --- | --- | --- | --- |
|  | 299.43 | 59.48 | 254.40 | 165.36 |
|  | 289.45 | 59.89 | 250.16 | 156.88 |
|  | 289.45 | 59.07 | 267.12 | 182.32 |
|  | 310.13 | 59.07 | 258.64 | 169.60 |
|  | 289.45 | 59.48 | 254.40 | 169.60 |
|  | 289.45 | 59.48 | 250.16 | 165.36 |
|  | 299.43 | 59.07 | 254.40 | 169.60 |
| **Mean** | 295.00 | 59.40 | 256.00 | 168.00 |
| **Standard Deviation** | 8.07 | 0.31 | 5.85 | 7.63 |
| **Standard Error** | 3.05 | 0.12 | 2.21 | 2.88 |

**Table 1. Measured parameters for individual Class I and Class II filaments.**
| **Class II** | **Crossover spacing** | **Turn layer-line** | **MT outer diameter** | **MT lumen diameter** | **Meridional** |
| --- | --- | --- | --- | --- | --- |
|  | 271.36 | 61.59 | 267.12 | 173.84 | 61.15 |
|  | 310.13 | 61.15 | 275.60 | 178.08 | 60.30 |
|  | 299.43 | 60.30 | 275.60 | 182.32 | 62.47 |
|  | 310.13 | 60.72 | 275.60 | 173.84 | 61.15 |
|  | 271.36 | 60.30 | 284.08 | 186.56 | 61.59 |
|  | 241.21 | 60.72 | 267.12 | 186.56 | 62.47 |
|  | 299.43 | 62.03 | 275.60 | 186.56 | 61.59 |
|  | 249.37 | 61.15 | 271.36 | 173.84 | 61.59 |
|  | 256.93 | 61.59 | 273.24 | 173.88 | 61.44 |
|  | 256.93 | 61.89 | 273.24 | 169.74 | 61.89 |
|  | 264.96 | 62.34 | 260.82 | 165.60 | 61.89 |
|  | 264.96 | 58.88 | 273.24 | 173.88 | 62.34 |
|  | 256.93 | 61.00 | 273.24 | 178.02 | 62.34 |
|  | 264.96 | 62.34 | 269.10 | 182.16 | 61.89 |
|  | 249.37 | 60.13 | 277.38 | 190.44 | 61.89 |
|  | 271.36 | 61.44 | 277.38 | 194.58 | 62.34 |
|  | 289.45 | 61.44 | 279.84 | 169.60 | 61.59 |
|  | 310.13 | 61.15 | 262.88 | 165.36 | 61.59 |
| **Mean** | 274.36 | 61.12 | 272.90 | 178.00 | 61.75 |
| **Std. Deviation** | 22.85 | 0.86 | 5.81 | 8.48 | 0.55 |
| **Std. Error of Mean** | 5.39 | 0.20 | 1.37 | 2.00 | 0.13 |

### Layer-line analysis confirms that lumenal filaments are composed of F-actin

To further characterise microtubule cores, lumenal regions were extracted from the tomographic reconstructions as 3D subvolumes. These were then summed in Z to obtain 2D projection images. Fourier transforms of these images were then calculated and inspected. Helical filaments like actin and tubulin display distinct layer line patterns. Data from single representative microtubules containing filaments of the Class I and Class II varieties are shown in Figure 3 and measured parameters from several microtubules/filaments are shown in Table 1. Class I filament containing microtubules display an actin layer line pattern - a clear reflection at 5.94 ± 0.03 nm - the pitch of the actin genetic helix and a reflection at 29.50 ± 0.80nm corresponding to the crossover spacing of the two long pitch actin helices. This crossover spacing is shorter than for canonical actin (~35nm) ^19^ but greater than that reported for actin cofilin filaments (~27nm) ^20^ and actin is known to have a random variable twist ^21^. These results enable us to confidently identify the Class I lumenal filament as F-actin. A reflection at 4nm was also visible from the tubulin monomer. Class II filament containing microtubules maintain an actin-layer line pattern with reflections at 6.11 ± 0.09 nm and 27.44 ± 2.29 nm, that is augmented by a strong reflection on the meridian at 6.18 ± 0.06 nm (matching the pitch of the genetic helix of actin). This meridional reflection is indicative an additional protein(s) associated with the lumenal actin. In Extended Data Figs. 5 and 6 we show that Class I actin filaments have symmetry close to an 11 subunit in 5 turn helix of axial repeat 29.5 nm, and that Class II actin filaments have symmetry close to a 20 subunit in 9 turn helix of axial repeat 2 x 27.5 nm. We tentatively speculate that the meridional reflection from Class II filaments is consistent with a formin-like encircling of the actin backbone ^22^.

## Discussion

The discovery of *m*icrotubule *l*umenal actin (ML-actin) in our cell-based extrusion system prompts the questions of how it is incorporated, its function(s) and abundance in settings that are not small-molecule modified. Two hypotheses present themselves that are not mutually exclusive. Firstly, altered microtubule dynamics caused by activation of kinesin-driven microtubule-microtubule sliding and bundling provides opportunity for actin to enter the microtubule lumen. This may occur through transient opening of the microtubule under strain when tight loops are formed (Extended Data. Fig. 1) or by kinesin-mediated disruption of the lattice ^9,23,24^. Put another way, F-actin incorporation may occur in response to mechanical or structural stress ^25^. Alternatively, microtubule extrusion has facilitated the analysis of a pool of pre-existing actin-containing microtubules with previously limited accessibility by high resolution cryo-EM. Indeed, the similarity of our images to the ‘dense core microtubules’ first observed in amphibian and rat neurons as well as platelets is striking ^1-3,26,27^. In either case, it will be important to understand if and how an F-actin core (of either class) alters the mechanical and dynamic properties the microtubule as well as explore this new concept as a new basis for actin-microtubule crosstalk in diverse settings ^28-31^.

## Acknowledgements

This work was supported by the Biotechnology and Biosciences Research Council (BBSRC) (BB/S000917/1) and a Lister Research Prize Fellowship to M.P.D.. D.M.P is supported by a British Heart Foundation Career Re-Entry Fellowship FS/14/18/3071), the Alan Turing Institute through a Turing Fellowship and the Academy of Medical Sciences by a Springboard award (SBF003\1142). We acknowledge access and support of the GW4 Facility for High-Resolution Electron Cryo-Microscopy, funded by the Wellcome Trust (202904/Z/16/Z and 206181/Z/17/Z) and BBSRC (BB/R000484/1) as well as the Wolfson Bioimaging Facility at Bristol. The Cryo-fluorescence microscope was supported by the BBSRC (BB/L014181/1). We are grateful to Professor Carolyn Moores (Birkbeck, University of London) for helpful discussions on the project and comments on the manuscript draft.

## Author contributions

Performed experiments – D.M.P, J.S., U.B., J.C., K.S., P.V., M.P.D.

Analysed data - D.M.P, J.S., P.V., J.S., P.V., M.P.D.

Wrote the draft – M.P.D.

All authors contributed to revision of the draft.

## Methods

### Cell culture and small-molecule treatment

HAP1 cells were obtained Horizon Discovery and cultured in Iscove’s Modified Dulbecco’s Medium (IMDM) containing 10% FCS and Penicillin/Streptomycin at 37ºC in 5% CO_2_ incubator. For fluorescence imaging, cells were plated onto fibronectin coated cover-slips in a 6-well plate at a density of 1×10^5^ per well the day before small-molecule treatment. Cells were prepared for electron microscopy as described below. Kinesore (3,5-dibromo-N′-[2,5-dimethyl-1-(3-nitrophenyl)-1H-pyrrol-3-yl]methylene}-4-hydroxybenzohydrazide) was obtained from Chembridge Corporation (Cat. No. 6233307) and prepared as a 50 mM stock in DMSO. In addition to manufacturer provided QC, molecular weight was verified by mass spectrometry. To stimulate projection formation. treatments were carried out in Ringer’s Buffer ([155 mM NaCl, 5 mM KCl, 2 mM CaCl_2_, 1 mM MgCl_2_, 2 mM NaH_2_PO_4_, 10 mM glucose, 10 mM Hepes (pH 6.8)] (in a 37ºC incubator without CO_2_) containing 100 µM Kinesore (final DMSO concentration 0.2%) for one hour. Control experiments were performed with DMSO alone in Ringers buffer at 0.2%.

### Live-cell imaging and analysis

To label tubulin, HAP1 cells were incubated with SiR-Tubulin (500nM) in growth media for 1 hour prior to kinesore treatment as described above, before transfer to the microscope stage. Images were acquired at 15 second intervals as single confocal sections, using a 633nm laser on a Leica SP5-II system with a 63x objective lens. Resulting data was further analysed as indicated and prepared for publication using Fiji (ImageJ) and Adobe Photoshop/Illustrator Packages.

### Correlative Light Electron Microscopy

Cells were grown on Quantifoil R1.2/1.3 400 mesh gold EM grids (supplied by EM Resolutions Ltd, UK). These were plunge frozen in liquid ethane using a Leica EM GP plunge freezer and transferred to a Leica CryoCLEM stage based on the Leica DM6000FS fluorescence light microscope. In this microscope the objective is cooled and the sample is transferred to a cryo stage where its temperature can be maintained below −140 °C during observation ensuring that the sample remains vitrified. Images were collected in both bright field and in green fluorescent channel. Areas of potential interest for further study by cryoTEM were thus recorded in a manner similar to ^32^. The samples were recovered under liquid nitrogen, transferred to a Tecnai20 LaB6 TEM (FEI), operating at 200kV using a Gatan 626 cryotransfer holder. The areas of interest were retraced and images collected on a bottom-mounted Thermo-Fisher CETA camera.

### Cryo Electron Tomography

Cryo samples were prepared as described above, clipped, and transferred into a Talos Arctica CryoTEM (FEI) operating at 200 kV. Bidirectional tilt series were collected with an angular range of +20° to −60° / +60°, angular increments of 3°, total dose of 109 e^−^/Å^2^, and a defocus range between −2 µm to −4 µm. Images were recorded in counted mode (2.21 Å/pixel) using a K2 Summit direct electron detector (Gatan) fitted to a BioQuantum energy loss spectrometer (Gatan) operating with a 20 eV slit width.

### Data processing

Reconstructions, movies and microtubule/actin filament length analysis were performed using IMOD (University of Colorado, Boulder)^33^ and Fiji (ImageJ) software.

### Layer Line analysis

Fourier transforms of 2D projections of extracted filament volumes were calculated and the layer-line positions measured using Fiji (ImageJ). The well characterized 4 nm reflection of the tubulin monomer was used as an internal calibration tool and the real space pixel size for each filament calculated. Modelling of the layer-line patterns from helices with actin-like symmetry (Supplementary Figures 5 and 6) was carried out using the HELIX program:

(Knupp & Squire: https://www.diamond.ac.uk/Instruments/Soft-Condensed-Matter/small-angle/SAXS-Software/CCP13/HELIX.html).

---

**Extended Data Fig. 1.**
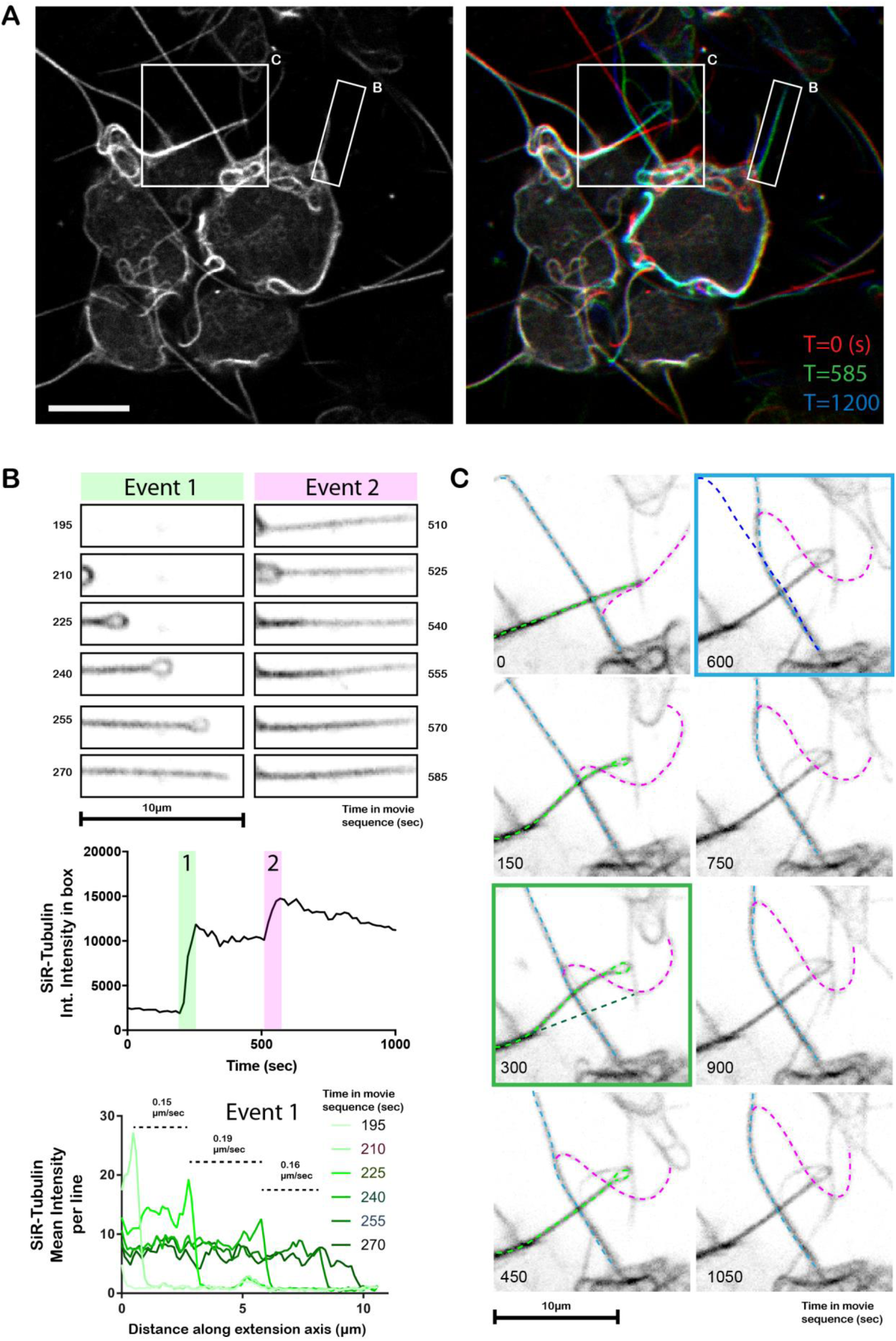
Live-cell imaging of microtubule projection formation through loop-extrusion. Live-cell confocal imaging of tubulin-labelled HAP1 cells (A) Left image shows cells at t=0 seconds. Right image shows T=0, T=585 and T=1200 (time in sec) in red, green and blue channels respectively. Boxes B and C highlight regions that are expanded and analysed in (B). and (C). Scale bar is 10 µm (B) Shows stills from the image series describing the formation and reinforcement of a microtubule-based projection through the extension/resolution of a loop. Green panels show extension of the first loop (event 1). Top graphs show change in total tubulin intensity over the whole region as a function of time. Lower graph shows average tubulin intensity per line as a function of distance along the axis of projection extension (x-axis) and rate of extension. Magenta panels (event 2) show addition of further tubulin intensity over a similar time period later in the movie, indicating that such projections are formed by the progressive layering of microtubules from the extrusion of loops. (C) Highlights complex interactions between filaments that affect their organisation. Three extensions are highlighted by light blue, light green and magenta dotted lines. The magenta projection moves along light blue projection. The position of the light green filament is shifted as it passes. Green box panel shows orginal (dark green) and new positions (light green) of the projection. As the magenta proceeds along the blue filament, the blue projection is bent. Blue boxed panel shows orginal (dark blue) and new positions (light blue) of the blue projection.

**Extended Data Fig. 2.**
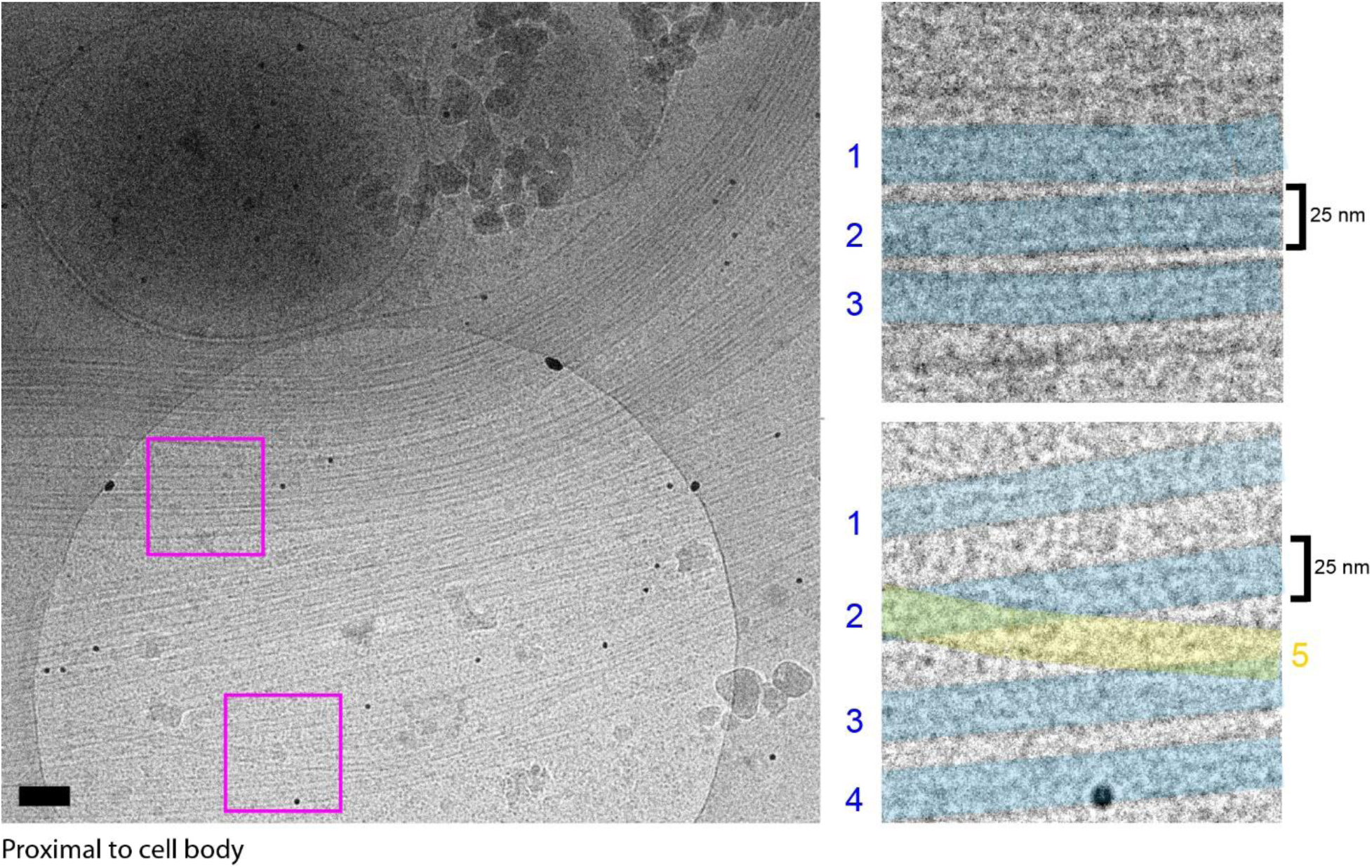
Cryo-CLEM reveals that projections are composed of closely aligned microtubule bundles. A single EM image acquired proximal to the cell body with a large number of microtubules. Boxes show microtubules with similar alignments (shaded blue) and an example of a single microtubule that crosses a bundle of aligned microtubules (yellow, 5). Scale bars 100 nm.

**Extended Data Fig. 3.**
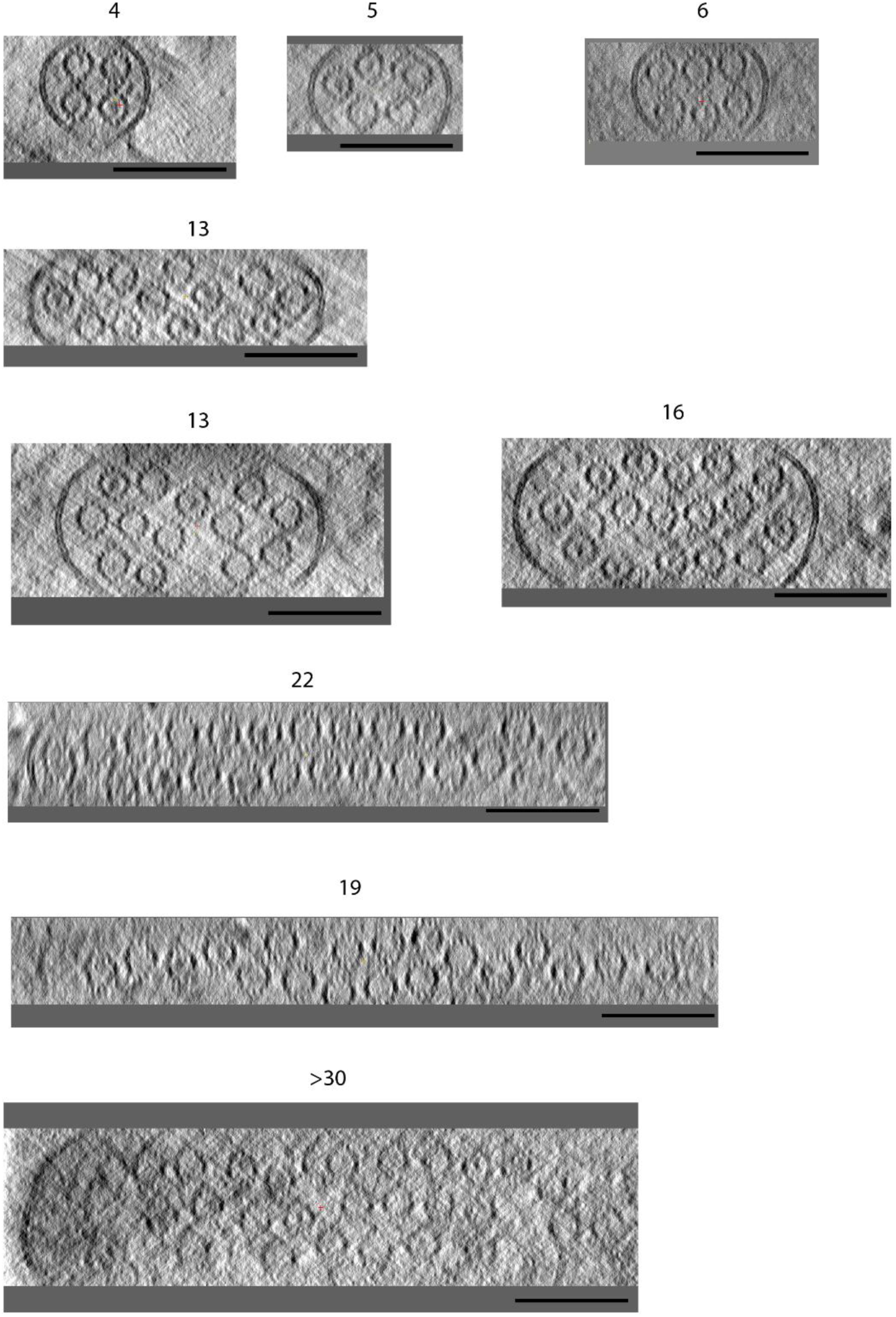
Sections through microtubule-based projections reveal variable microtubule number and close-packed organisation. Transverse sections through several microtubule-based projections show the close juxtapositon of variable numbers of microtubules. Lumenal density in many of the microtubules is apparent. Images are presented as summed slices. Scale bars 100 nm.

**Extended Data Fig. 4.**
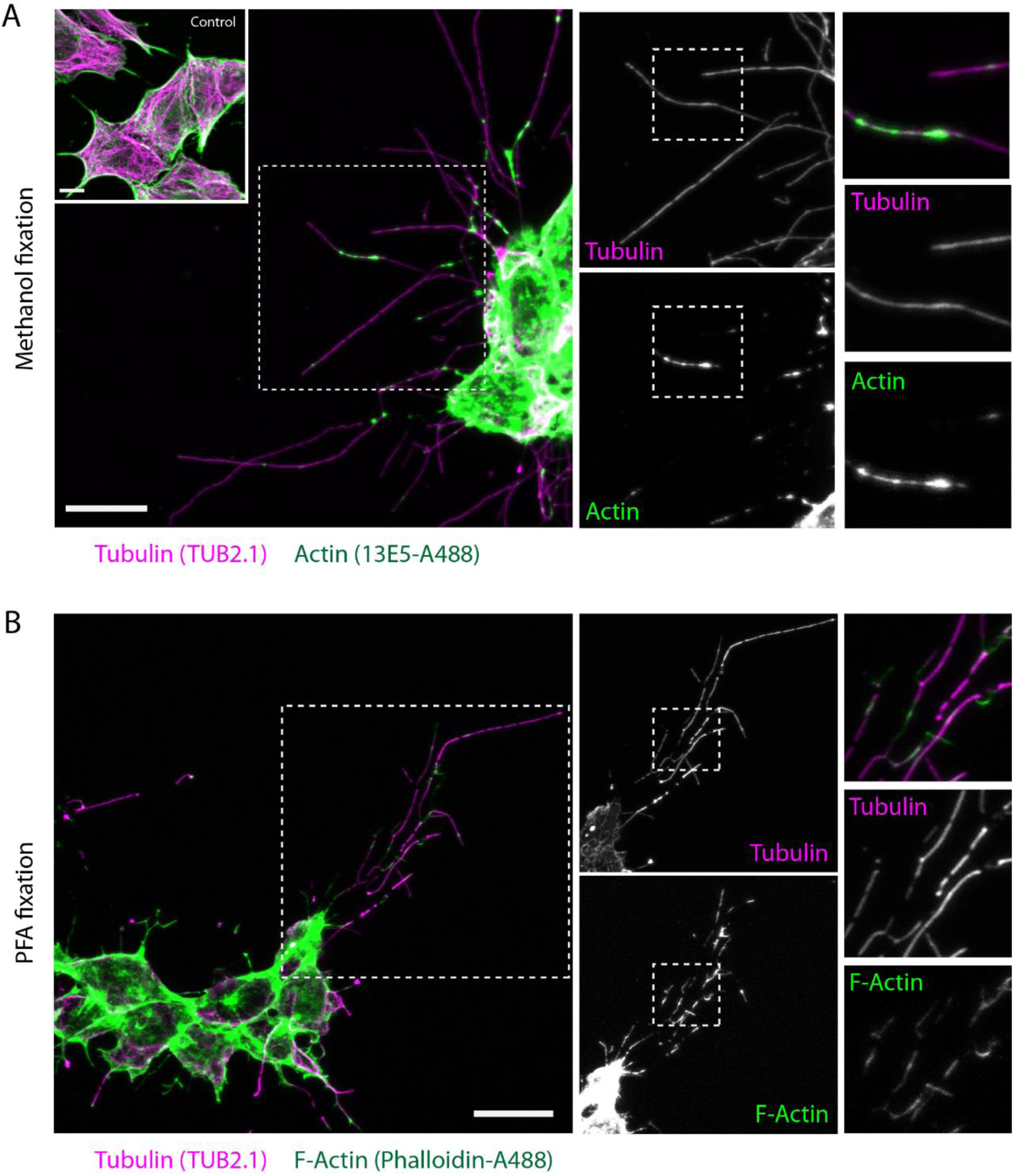
Kinesore-induced projections contain both tubulin and F-actin. HAP1 cells were treated with kinesore (100µM) for one hour and fixed with either ice cold methanol (A) or 4% paraformaldehyde (B). (A) Methanol fixed cells were stained with antibodies against tubulin (TUB 2.1, detected by anti-mouse Alexa Fluor 568) or actin (Rabbit anti β-actin (1E35) directly conjugated with Alexa Fluor 488). Inset shows control cells treated with buffer/vehicle only (B) PFA fixed cells were stained with TUB2.1 and Phalloidin-Alexa Fluor 488 to detect filamentous actin (F-actin). Boxed regions show cellular projections at increasing magnification and highlight overlapping patterns in actin and tubulin staining. Scale bars are 10 µm.

**Extended Data Fig. 5.**
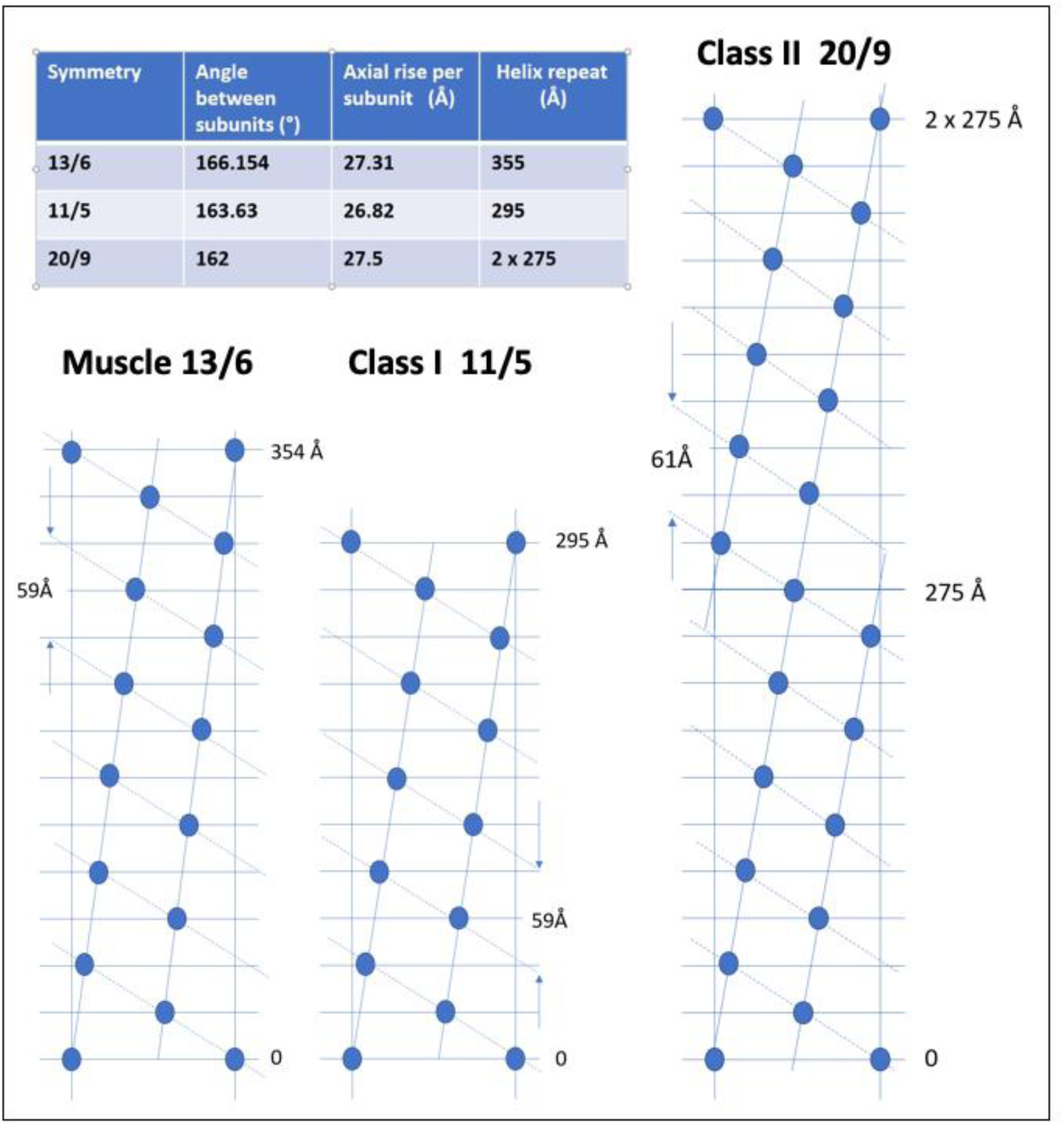
Actin filament radial nets. Comparison of the radial nets of muscle actin filaments (13/6 helices of actin monomers) and Class I and Class II actin filaments as observed here, where Class I filaments have approximate 11/5 helical symmetry and Class II filaments have 20/9 helical symmetry. The inset Table shows the helical parameters for the three symmetries.

**Extended Data Fig. 6.**
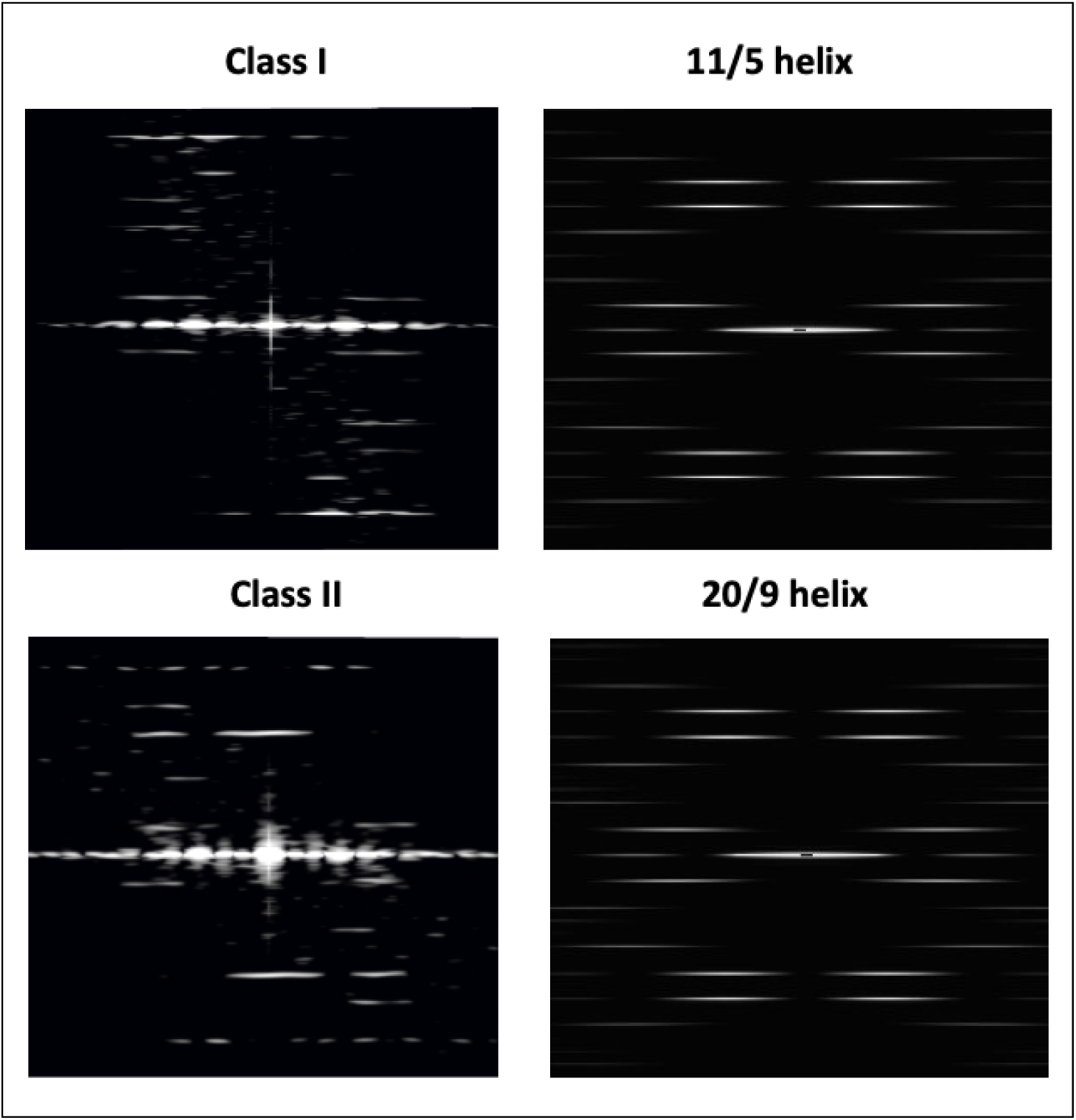
Diffraction from Class I and Class II filaments. The images on the left reproduce the patterns shown in main text Figure 3 for Class I (top) and Class II (bottom) filaments. On the right are patterns produced by the HELIX program (Knupp and Squire) for actin-like filaments with 11/5 (top) and 20/9 (bottom) helical symmetry. A filament length of 100 subunits (simple spheres) was used with the monomer at 25 Å from the filament axis

## Supplementary Videos

### Supplementary Movie 1

Movie shows HAP1 cells labelled with SiR-tubulin and treated with kinesore that is also shown in Supplementary Figure 1. Images were acquired every 15 seconds. Boxed region shows area of movie described in more detail in figure 1.

### Supplementary Movie 2

Movie shows a series of images through the tomogram described in detail in Figure 1B. Microtubules in the tomogram are then identified by coloured cylinders.

### Supplementary Movie 3

Movie shows Z-series through a subvolume containing two microtubules. The top microtubule contains a Class I filament, the bottom microtubule contains globular density

